# Forest expansion and open vegetation responses during the past 14 ka at Zhagaer Co on the eastern Tibetan Plateau

**DOI:** 10.64898/2026.04.29.721606

**Authors:** Yanrong Zhang, Xianyong Cao, Kathleen R. Stoof-Leichsenring, Shengqian Chen, Fang Tian, Ulrike Herzschuh

**Affiliations:** Alfred Wegener Institute, Helmholtz Centre for Polar and Marine Research, Polar Terrestrial Environmental Systems, Potsdam 14473, Germany; Institute of Environmental Science and Geography, University of Potsdam, Potsdam 14476, Germany; Alpine Paleoecology and Human Adaptation Group (ALPHA Group), State Key Laboratory of Tibetan Plateau Earth System, Environment and Resources, Institute of Tibetan Plateau Research, Chinese Academy of Sciences, Beijing 100101, China; College of Resource Environment and Tourism, Capital Normal University, Beijing 100048, China; Institute of Biochemistry and Biology, University of Potsdam, Potsdam 14476, Germany

**Keywords:** sedaDNA, pollen, forest-alpine transitions, ecological filtering, Holocene vegetation change, alpine plant communities

## Abstract

Forest expansion and retreat are key drivers of alpine ecosystem dynamics, yet it remains unclear how forest expansion shapes alpine plant communities and whether alpine assemblages that re-emerge after forest decline resemble those of the Late Glacial. Here, we integrate sedimentary ancient DNA (sedaDNA) and pollen records from Zhagaer Co on the eastern Tibetan Plateau to reconstruct vegetation changes over the past ∼14 ka. Using *Picea* abundance as a proxy for forest cover, we apply constrained ordination of sedaDNA-inferred plant communities to identify plant taxa as “winners” and “losers” of forest expansion and compare the composition of loser taxa between the late Glacial and late Holocene cold-open phases. Our results show that forest expansion during the early to mid-Holocene favoured woody and forest-margin taxa (e.g. *Rhododendron*, Salicaceae), while suppressing alpine forbs and graminoids (e.g. *Carex*, *Thalictrum*), consistent with patterns expected under ecological filtering. However, late Holocene reopening did not fully restore a late Glacial-like alpine community; instead, it was characterised by a stronger contribution of alpine meadow and shrub taxa. This difference may reflect contrasting environmental backgrounds, including higher atmospheric CO2 levels during the late Holocene, ecological legacies of prior forest expansion, and increasing human influence. These findings suggest that forest expansion may lead to long-term restructuring of alpine plant assemblages, and that late Holocene cooling did not simply restore late Glacial alpine communities but instead produced a distinct alpine ecosystem state. Together, these results highlight the long-term legacy of treeline dynamics in shaping alpine ecosystem trajectories.

## 1. Introduction

Forest–alpine ecotones are among the most climate-sensitive vegetation boundaries, where treeline position and ecotone structure reflect interactions between climatic constraints and local processes (Holtmeier & Broll, 2020; Bader et al., 2021). Across high mountain systems, Holocene forest expansion and retreat have been widely documented through pollen records and other palaeoecological archives, demonstrating substantial shifts in treeline position and woody cover in response to climatic forcing (Schwörer et al., 2017; Marquer et al., 2020). These transitions are commonly interpreted as temperature-driven changes in habitat extremes (Harsch et al., 2009), with shifts in alpine habitat area assumed to alter community composition (e.g. Liu et al., 2021). However, relatively few studies have explicitly evaluated whether alpine plant communities that re-establish following forest retreat are compositionally comparable to pre-expansion alpine states. In particular, it remains unclear whether complete forest–alpine cycles result in reversible changes in community composition, in terms of both taxonomic structure and relative dominance.

Forest expansion is widely recognized as a key driver of ecological change in alpine ecosystems, affecting habitat availability, plant diversity and community composition (Harsch et al., 2009; Kumar & Khanduri, 2024). However, much of our current understanding derives from studies documenting tree encroachment and forest advance over relatively short observational timescales (Greenwood & Jump, 2014; Holtmeier & Broll, 2020). Consequently, the ecological consequences of forest retreat following earlier expansion remain poorly understood. Long-term palaeoecological records provide a unique opportunity to examine complete forest–alpine cycles encompassing both forest expansion and retreat, and to compare alpine communities that develop under broadly similar open conditions, but under different climatic regimes and historical contexts. This framework allows us to assess whether alpine communities that re-establish following forest retreat resemble their pre-expansion states, or instead reflect persistent changes in community composition.

The late Glacial and the late Holocene both encompass cold episodes that promoted expanded open-alpine conditions in mountain systems, and may thus be regarded as broadly comparable cold–open states, despite differences in magnitude, temporal pacing, and atmospheric CO2 concentrations, which may also influence vegetation dynamics (Wanner et al., 2008; Rasmussen et al., 2014; Herzschuh et al., 2011). Pollen records from the Tibetan Plateau and surrounding regions commonly show that both periods are dominated by open vegetation, giving the impression of broadly similar community compositions (Herzschuh et al., 2014). However, a crucial difference between these two periods is that the late Holocene followed a prolonged phase of forest expansion during the early to mid-Holocene, when alpine habitats were compressed and potentially subjected to trait-based ecological filtering (Körner, 2012; Suding et al., 2008). Prolonged forest filtering may have reduced the representation of light-demanding alpine taxa, thereby altering the composition of subsequent late Holocene cold–open communities (Cao et al., 2013; Suding et al., 2008). This raises the question of whether the apparent similarity suggested by pollen records reflects true compositional equivalence, or whether higher-resolution data would reveal distinct ecological states shaped by legacy effects of prior forest expansion. This contrast therefore provides a natural framework to test whether alpine ecosystems show compositional equivalence under broadly comparable climatic conditions, or whether prior forest expansion imposes a lasting trajectory dependence on community assembly (Herzschuh, 2020; Johnstone et al., 2016).

Here, we analyse sedaDNA and pollen records from Zhagaer Co on the eastern Tibetan Plateau to examine vegetation dynamics across late Glacial–Holocene forest expansion and subsequent reopening. Using a forest-cover gradient represented by *Picea*, a dominant treeline-forming taxon and a key indicator of forest expansion in the study region (Schlütz & Lehmkuhl, 2009), in redundancy analysis (RDA), we identify taxa associated with forest-dominated and open-alpine conditions and characterise relative winner–loser dynamics. We then compare taxa aligned with open conditions during the late Glacial and the late Holocene to evaluate whether communities reassemble along similar trajectories following treeline retreat. We test two hypotheses: (1) taxa associated with woody and forest-margin habitats increase along the forest-cover gradient, whereas light-demanding alpine specialists decline; and (2) If forest expansion imposes ecological filtering on alpine communities, taxa that decline during forest expansion will show altered representation following late Holocene treeline retreat, indicating trajectory dependence in community assembly.

## 2. Study area

Zhagaer Co (33.064328°N, 101.17601°E, 4220 m a.s.l (above sea level)) is located in the southeastern part of Nianbaoyuze, eastern of Tibetan Plateau (Figure 1a, 1b). It lies in Sichuan province, in Aba Tibetan and Qiang Autonomous Prefecture of China. The Nianbaoyuze massif rises about 500–800 m above the surrounding landscape, and late Pleistocene glaciation left numerous glacial landforms and lakes in the region (Lehmkuhl and Liu, 1994; Lehmkuhl, 1995), including Zhagaer Co. The lake, encompassing an area of approximately 0.3 km², exhibits a maximum depth exceeding 30 meters, an exceptionally low pH value of 0.001, and is characterized as a glacial freshwater lake. It is enclosed on three sides by steep, towering mountains with an average relative relief of about 700 meters. It is primarily fed by snowmelt and local precipitation, with a river outlet located along the northern margin of the lake. The study area is influenced by the East Asian and South Asian monsoons. The mean annual temperature is 6.73°C and the mean annual precipitation is 705.17 mm, with precipitation primarily concentrated in the summer months (May to September) and the data from the nearest meteorological station to Zhagaer Co, Aba County (32.9°N, 101.7°E, 3275.1m a.s.l, ∼51 km from the lake), for the period 1954–2008 CE (Figure 1e, National Meteorological Information Center, 2019).

**Figure 1.**
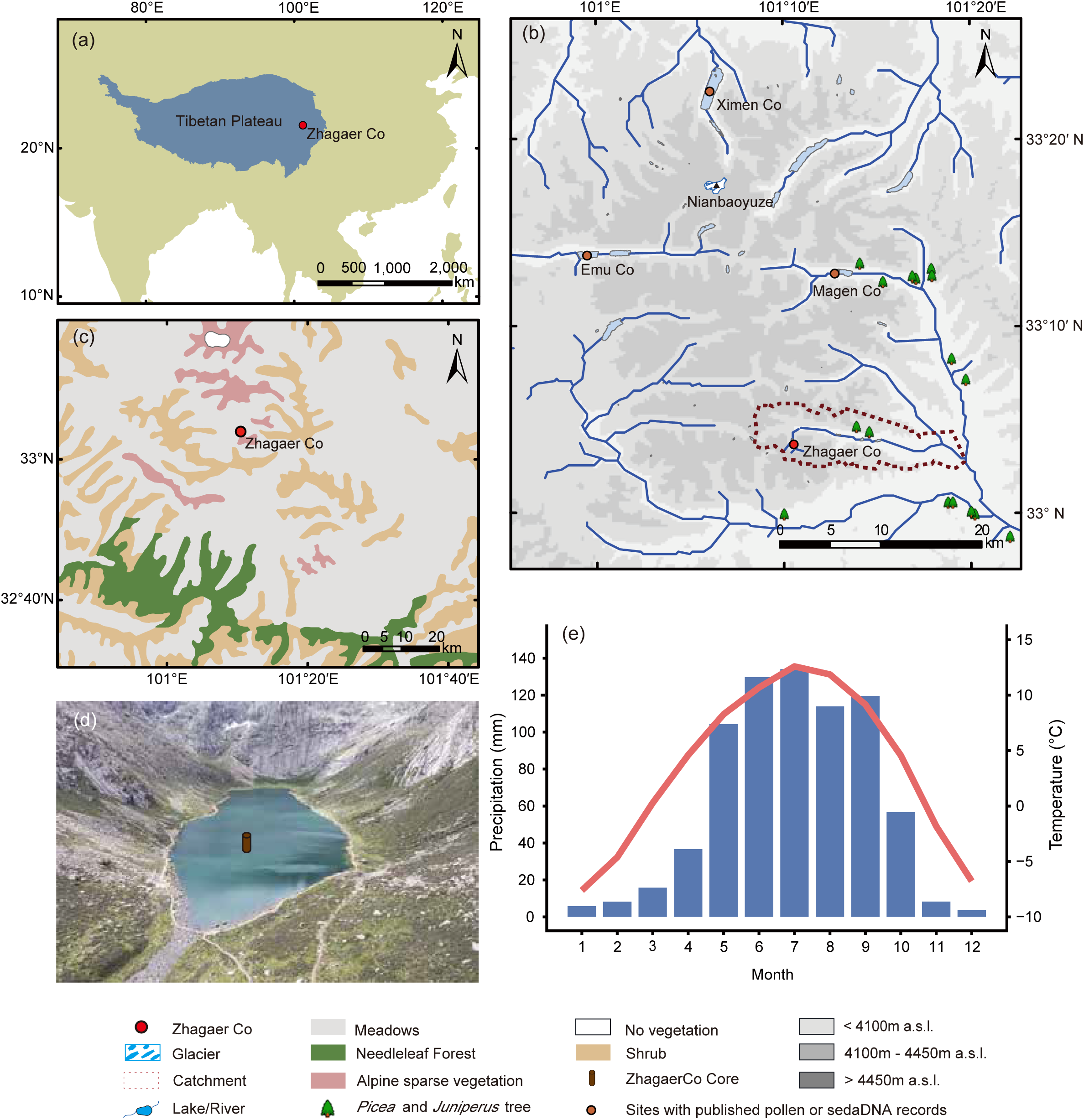
Location and environmental setting of Lake Zhagaer Co. (a) Location of the Tibetan Plateau; (b) map of the Nianbaoyeze Mountains showing the location of Zhagaer Co and other nearby lakes with published pollen or sedaDNA records; (c) location of Zhagaer Co and the type of vegetation around it; (d) location of the core in Zhagaer Co; (e) Climatology of Aba county (1954–2008 CE)

Zhagaer Co is located at high elevation, where vegetation shows pronounced vertical zonation along topographic and altitudinal gradients (Figure 1c). Within the lake catchment, the lower reaches of the basin and valley bottoms have arboreal vegetation dominated by cold-tolerant conifers, with *Picea* as a characteristic taxon. With increasing elevation, tree cover declines and vegetation transitions to shrub-dominated communities. Shrubs are mainly represented by Ericaceae (e.g., *Rhododendron bamaense*, *R. nitidulum*), Salicaceae (e.g., *Salix oritrepha*), and Rosaceae (e.g., *Spiraea alpina*, *Dasiphora parvifolia*). In areas above the lake, the vegetation is mainly composed of alpine taxa, such as Cyperaceae (*Carex capillifolia*, *Carex montis-everesti*), Polygonaceae (*Bistorta vivipara*), and Asteraceae (*Taraxacum calanthodium*). Around Zhagaer Co, vegetation is likewise dominated by shrubs and forms alpine shrub meadow and alpine meadow communities, in which Ericaceae shrubs are prominent and are accompanied by alpine herbaceous taxa, including Papaveraceae (*Meconopsis*), Saxifragaceae. Further upslope in the upper catchment, vegetation is mainly characterised by alpine meadows including Polygonaceae (*Polygonum*), Cyperaceae (*Kobresia*), Ranunculaceae (*Anemone*), Rosaceae (*Rubus*).

## 3. Materials and methods

### 3.1 Sediment core and chronology

In June 2023, a 740-cm-long sediment core (ZGEC) was obtained from the center of Zhagaer Co at a water depth of 29.95m (Figure 1d) using the UWITEC piston. The pipe for storing sediment is 3 m in length and the sample is 680 cm after deducting the overlapped part. A 76-cm gravity sediment was collected by UWITEC gravity corer at a location 1 meter away from the long sediment core. After field work, the core was promptly transported to the State Key Laboratory of Tibetan Plateau Earth System and Resources Environment, China. A total of fourteen bulk organic carbon were selected for accelerator mass spectrometer (AMS) 14C dating at Guangzhou Carbon Age Analytical Laboratory Co., Ltd. (Guangzhou, China) and at the MICADAS (Mini Carbon Dating System) Laboratory of AWI in Bremerhaven (Mollenhauer et al., 2021). Sediment samples were taken from different core depths and used for AMS 14C dating (Table 1). Three samples from the 60–160 cm depth interval yielded anomalous radiocarbon ages. Additional samples from the corresponding stratigraphic intervals are currently being selected for AMS 14C dating in order to refine the age–depth model. The determined lake reservoir effect is 256 years. The results will be incorporated into the final age–depth model once the new measurements are available.

**Table 1.** AMS 14C dating of bulk sediment from the Zhagaer Co sediment core.

| Lab ID | Depth (cm) | 14C age (year BP) | Error (years) |
| --- | --- | --- | --- |
| AWI13240 | 1 | 183 | 22 |
| AWI13241 | 21 | 1333 | 22 |
| TN20294 | 244 | 5385 | 30 |
| AWI13243 | 291 | 5999 | 25 |
| TN20295 | 344 | 7170 | 35 |
| AWI13244 | 391 | 7817 | 28 |
| TN20296 | 444 | 9275 | 50 |
| TN20297 | 489 | 9370 | 45 |
| AWI13245 | 541 | 11066 | 37 |
| TN20298 | 589 | 11190 | 60 |
| AWI13246 | 671 | 11938 | 42 |

### 3.2 sedaDNA analyses

In November 2023, 68 subsamples were taken from Zhagaer Co in a climate chamber maintained at the German Research Centre for Geosciences Helmholtz Centre Potsdam at a temperature of 4°C. Prior to core processing, all surfaces in the climate chamber were cleaned with DNA Exitus Plus™ (VWR, Germany) followed by demineralised water. Sampling tools, such as knives, scalpels, and their holders, were cleaned before each use and about 2mm of the exposed sediment of samples was removed with a small single-use clean blade and the inner part used for ancient DNA isolation, following the clean split sampling protocol described by following the procedure described by Parducci et al. (2017) and von Hippel et al. (2022). Collected sediment samples were stored in 8 mL sterile tubes (Sarstedt) at −20 °C.

All DNA extraction, amplification, and purification work were carried out in ancient DNA dedicated facilities at Alfred Wegener Institute, Helmholtz Centre for Polar and Marine Research Potsdam, using strict ancient DNA precautions and protocols (Zimmermann et al., 2017). Each extraction batch included nine samples (around 3g sample^−1^) and one extraction control, which was treated with a partially modified protocol of PowerMax^Ⓡ^ Soil DNA Isolation kit (Qiagen). DNA extracts from the samples were purified and concentrated using the GeneJET PCR PurificationKit (Thermo Scientific). PCR (Polymerase chain reactions) was performed with the “g” and “h” universal plant primers for the P6 loop region of the chloroplast *trn*L (UAA) intron (Taberlet et al., 2007). All primers were modified with an NNN-8 bp tag to establish sample multiplexing. PCRs were run in three replicates along with No Template Controls (NTCs) to control chemical contamination of PCR chemicals. PCR products were amplified in a Biometra ThermoCycler (Biometra, Germany) at initial denaturation at 95°C for 5 min, followed by 40 PCR cycles with denaturation at 95 °C for 30 s, annealing at 50 °C for 30 s and elongation at 68 °C for 30 s. Each PCR product has been checked by gel electrophoresis using a2% agarose gel (Jena Analytics). Purification of PCR products for each sample was done with MinElute (Qiagen, Germany), following the manufacturer’s protocol and 3×88 PCR products were pooled in equimolar concentrations. DNA concentrations of DNA extracts and purified PCR products were measured with the Qubit 4.0 fluorometer (Invitrogen), and the results showed no obvious contamination during the laboratory work. PCR-free library preparation and next-generation sequencing (2 × 150 bp) on the Illumina NextSeq 500 platform were performed at Genesupport LifeScience Fasteris SA in Switzerland.

Raw sequencing data were processed using the OBITools3 pipeline (Boyer et al., 2016). Paired-end reads were merged using obi alignpairedend, demultiplexed by tag combinations with obi ngsfilter, and dereplicated using obi uniq. PCR and sequencing errors were filtered using obi clean. Taxonomic assignment was performed with obi ecotag against a customized Tibetan plant (“TP_db”) reference database (Li et al., under review).

After processing the data using OBITools3, ASVs with less than 100% taxonomic identity were excluded. In addition, to restrict the analyses to terrestrial plant communities, all ASVs assigned to aquatic taxa were removed prior to downstream analyses. Non-metric multidimensional scaling (NMDS) analysis was subsequently performed via the metaMDS function in the Rpackage “*vegan*” to check the PCR replicability of the samples.

### 3.3 pollen analyses

Pollen samples were taken from the same 68 sediment levels used for sedaDNA analysis. Due to poor pollen preservation at one level (depth 111 cm), only 67 samples could be reliably identified and included in subsequent analyses. The samples generally weight 1–2 g and were pre-treated using standard laboratory methods (Fægri et al., 1989), including 10% HCL-10% NaOH-40% HF to remove carbonates, silicates, and humic acid in the sample and acetolysis (9:1 mixture of acetic anhydride and sulphuric acid), followed by a 7 μm mesh sieving and mounting in silicone oil. Before these treatments, *exotic Lycopodium spores* (10,315 spores per tablet) were added to each sample prior to processing to allow standardisation of pollen counts. Pollen identification was made with the help of published atlases for pollen (Wang et al., 1995; Tang et al., 2016; Cao et al., 2020). >500 terrestrial pollen grains were counted for pollen samples. Pollen percentage diagrams were produced in R using the function *strat.plot* in the rioja package (Juggins, 2023), which generates stratigraphic plots comparable to traditional Tilia-style pollen diagrams. Stratigraphic zonation was determined using CONISS clustering (Grimm, 1987) implemented in the same package.

### 3.4 Statistical analyses

All statistical analyses were performed in R. Prior to ordination analyses, rare and sporadically occurring taxa were filtered to reduce noise and improve the robustness of multivariate analyses. Taxa were retained only if they met both of the following criteria: (i) occurrence in at least three samples, and (ii) a maximum relative abundance of at least 0.1% in at least one sample.

We used RDA to quantify how plant community composition covaried with the abundance of *Picea* at Zhagaer Co. *Picea* was used as a proxy for forest expansion because it is a modern dominant tree taxon in the mountains of the eastern Tibetan Plateau (Cao et al., 2024). Modern vegetation surveys indicate that Picea forests occur in the downstream catchment of Zhagaer Co (Schlütz & Lehmkuhl, 2009), suggesting that increases in *Picea* abundance in the *seda*DNA record likely reflect regional forest expansion. The analysis was based on a terrestrial plant sedaDNA abundance matrix, with sample age retained as metadata and used solely for visualisation. Prior to ordination, taxon abundances were transformed using a fourth-root transformation to down-weight dominant taxa and improve ordination stability. Because *Picea* served as the constraining variable, its abundance was extracted from the community matrix and used as the sole explanatory variable, while the remaining taxa constituted the response matrix. RDA was performed using the rda function in the vegan R package, with *Picea* treated as a continuous predictor (model: community ∼ *Picea*). Site scores and biplot scores for the constraining variable were extracted using scores with scaling = 2. Samples were visualised in constrained ordination space in a site-only biplot, with the *Picea* variable displayed as a biplot arrow and axis labels indicating the proportion of variance explained. To aid interpretation of sample distribution, samples were classified into four quadrants based on the signs of their scores on the first two ordination axes.

To visualise how individual taxa were distributed along the *Picea* constraint while avoiding overplotting, we generated a taxon-only RDA biplot based on the same constrained ordination model (community ∼ *Picea*). Taxon scores and biplot scores for the constraining variable were extracted using scores in vegan with scaling = 2. Each taxon was assigned to one of four quadrants (Q1–Q4) based on the sign of its scores on the first two ordination axes. To quantify the strength of taxon separation in ordination space, we calculated vector length as the Euclidean distance from the origin (len = √(x² + y²)). For clarity, only taxa with the longest vectors (top 30 taxa ranked by *len*) were labelled in the biplot, while all taxa were retained in the ordination model. The direction of the *Picea* constraint was displayed as a biplot arrow, and axis labels indicate the percentage of explained variance. Quadrant-specific taxa lists were summarised and placed as tables in the corresponding corners of the ordination plot (Q1–Q4).

## 4. Results

### 4.1 Chronology

The radiocarbon age of the 0–1 cm surface sample indicates an apparent surface-age offset of 256 years relative to the sampling year, which was applied as the reservoir correction in the age model. In total, 11 samples from the sediment core were selected for AMS 14C dating, including the 0–1 cm surface sample used to estimate the reservoir offset. We then used the “Bacon” package, incorporating 14C ages and the reservoir correction, to construct the chronology for the sediment core (Figure 2). This chronology indicates that the sediment core spans the past 13.7 cal ka BP.

**Figure 2.**
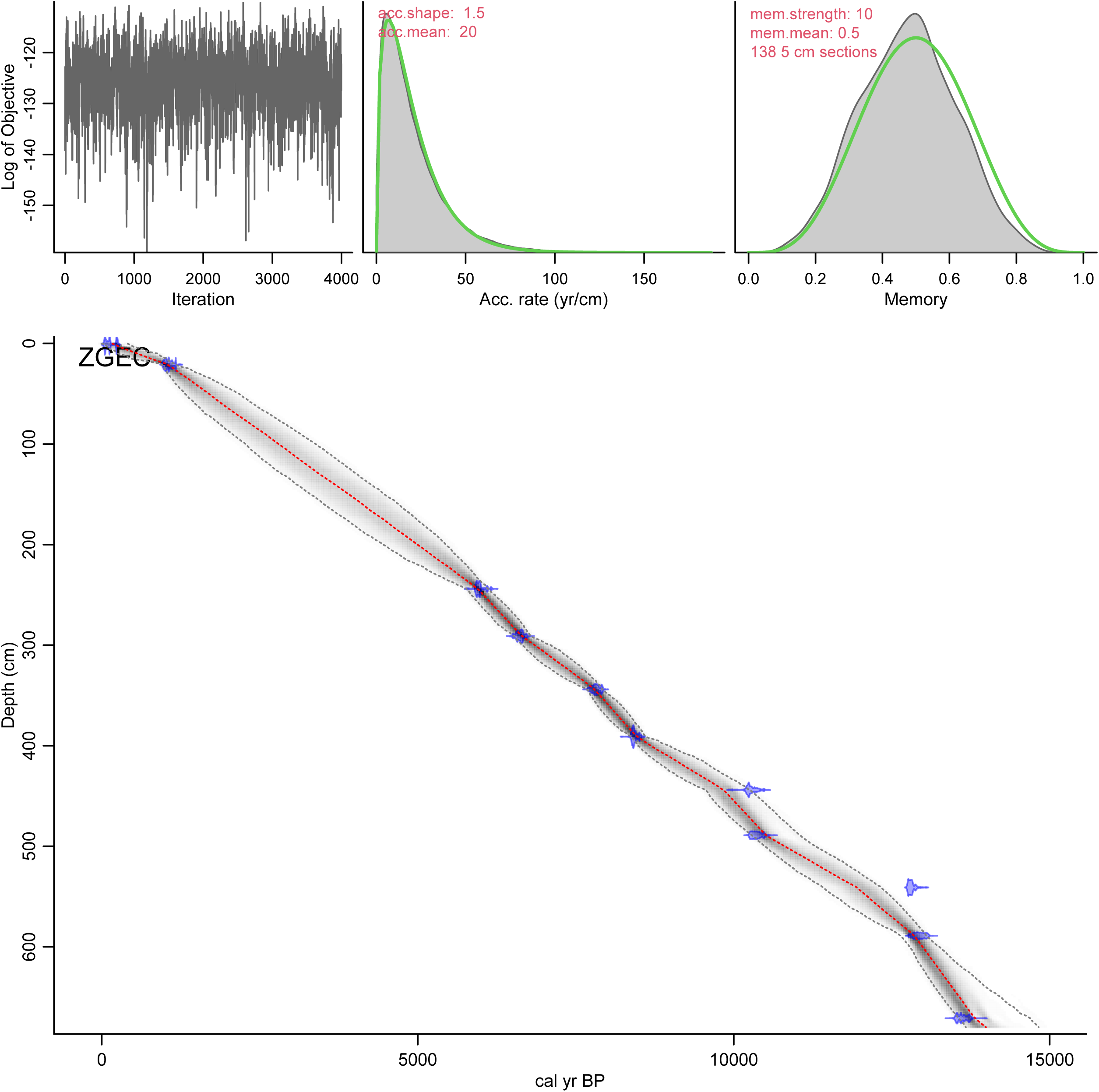
Age-depth relationship for the sediments of Zhagaer Co using Bacon model.

### 4.2 Plant DNA metabarcoding

In total, 546 ASVs have 100% sequence similarity with the “TP_db” database. Of these, 513 sequences are assigned to terrestrial seed plant taxa, and 48.3%, 34.1%, and 4.09% of them are identified to species, genus and family level, respectively. NMDS results show high similarity in taxonomic composition among the three PCR replicates of each sample, except for slight variations in the top two samples (Additional details are provided in the Supplementary Note 1).

Zonation was implemented using the CONISS method (chclust procedure), and clustering of relative-abundance data identified three major stratigraphic zones, reflecting pronounced shifts in terrestrial plant community composition through time (Figure 3). Zone I (late Glacial: 681–531 cm; 13,745–11,660 cal yr BP): Plant metabarcoding assemblages in Zone I are dominated by herbaceous alpine taxa, with frequent occurrences of *Ranunculus nephelogenes*, *R. rufosepalus*, *R. bungei*, *Meconopsis integrifolia*, *Micranthes divaricata*, *Saxifraga sinomontana*, *Saussurea dzeurensis*, *Carex przewalskii*, *Bistorta macrophylla*, Pooideae, *Astragalus*, and *Pedicularis*. Woody taxa are generally rare throughout this interval. Tree taxa are largely absent, although *Alnus cremastogyne* occurs at low abundance. Among shrubs, Saliceae are consistently present and locally reach moderate abundances, whereas other shrub taxa (e.g., *Lonicera*, *Hippophae tibetana*, and *Potentilla*) occur at lower levels; Zone II (early to mid-Holocene: 531–211 cm; 11,660–5,241 cal yr BP) is characterised by an expansion of arboreal taxa, notably *Picea*, Cupressaceae, *Juniperus microsperma*, and *Betula*. This increase in tree taxa is accompanied by higher representation of shrub taxa, including *Rhododendron*, *Rhododendron leptothrium*, Salicaceae, and *Ribes*. Although the overall contribution of herbaceous taxa declines relative to Zone I, changes within the herb layer are heterogeneous. Several alpine- or open-habitat–associated forbs, such as *Saussurea* (including *S. mucronulata*), Apioideae, and *Urtica*, appear or increase during this interval. Zone III (late Holocene: ca. 221–0 cm; 5,241 cal yr BP to present) is characterised by reduced arboreal representation and a renewed dominance of shrubs, forbs, and graminoids. Shrub taxa, including *Rhododendron leptothrium*, Rosoideae, Potentilleae, and *Lonicera*, increase during this interval, while the herbaceous component becomes more diverse and abundant. Prominent forbs include *Thalictrum*, *Silene*, *Delphinium*, Anthemideae, *Corydalis*, *Koenigia*, *Saxifraga*, *Chrysosplenium axillare*, *Ranunculus*, and *Caltha* (e.g., *C. scaposa*). Graminoids such as *Carex*, Poeae, and Pooideae are also well represented. In addition, several alpine-associated taxa increase or reappear during this interval, including *Saxifraga*, Apioideae, *Meconopsis*, *Gentiana*, *Ranunculus*, and *Rheum*.

**Figure 3.**
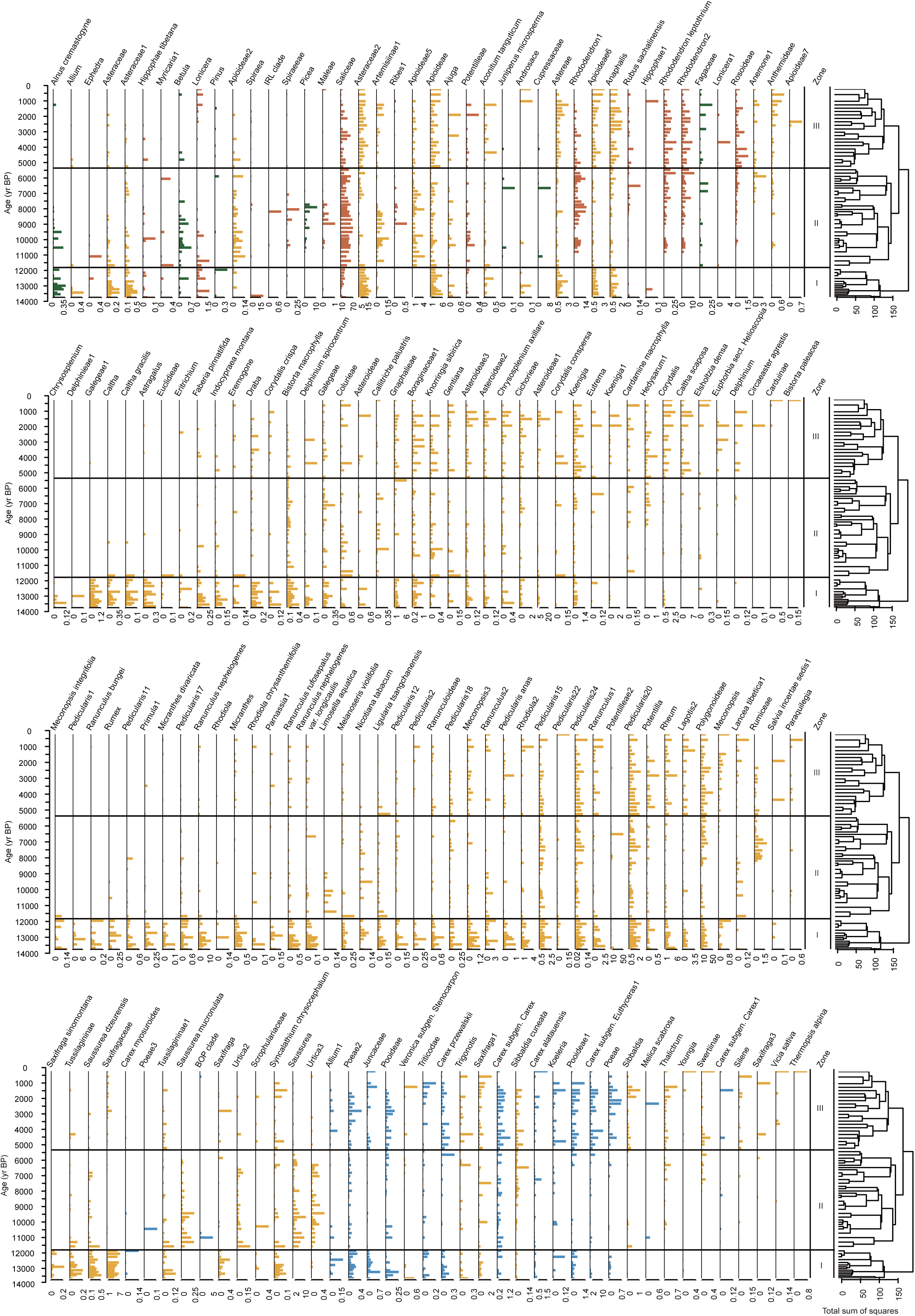
Relative abundance of terrestrial sedaDNA taxa over the past 14 ka from the Zhagaer Co sediment core. Green denotes trees, brown shrubs, yellow forbs, and blue graminoids. CONISS-defined zones are indicated on the right.

### 4.3 Pollen assemblages

A total of 56 pollen and spore types were identified from the Zhagaer Co sediment core, including 16 arboreal taxa, 13 shrub taxa, 16 herbaceous taxa, 2 hydrophyte taxa, and 4 algae and spore taxa. The herbaceous taxa (44.84–92.59%, mean = 75.22%), dominated by Cyperaceae (up to 58.92%), Poaceae (up to 32.74%) and Artemisia (up to 31.68%). Arboreal pollen abundance ranged from 3.04% to 50.96%, and main taxa are *Picea* (up to 29.33%), *Pinus* (up to 6.61%), *Betula* (up to 29.83%).

Three pollen zones were identified by CONISS analysis based on the abundance changes in pollen taxa (Figure 4). Pollen Zone I (late Glacial: 681–531 cm; 13,745–11,660 cal yr BP) is mainly characterized by a high abundance of herbaceous taxa, with high abundances of Cyperaceae, Poaceae and *Artemisia*. Pollen Zone II (early to mid-Holocene: 531–241 cm; 11,660–5,895 cal yr BP), the abundance of arboreal pollen increases, with *Picea*, *Betula*, *Pinus*. Pollen Zone III (241–0 cm; 5,895–0 cal yr BP), with the decreased main arboreal pollen abundance, and Cyperaceae, Poaceae and *Artemisia* are again dominated compared with Zone II. The alpine families, Saxifragaceae and Ranunculaceae, are also having higher abundance during Zone I and Zone III.

**Figure 4.**
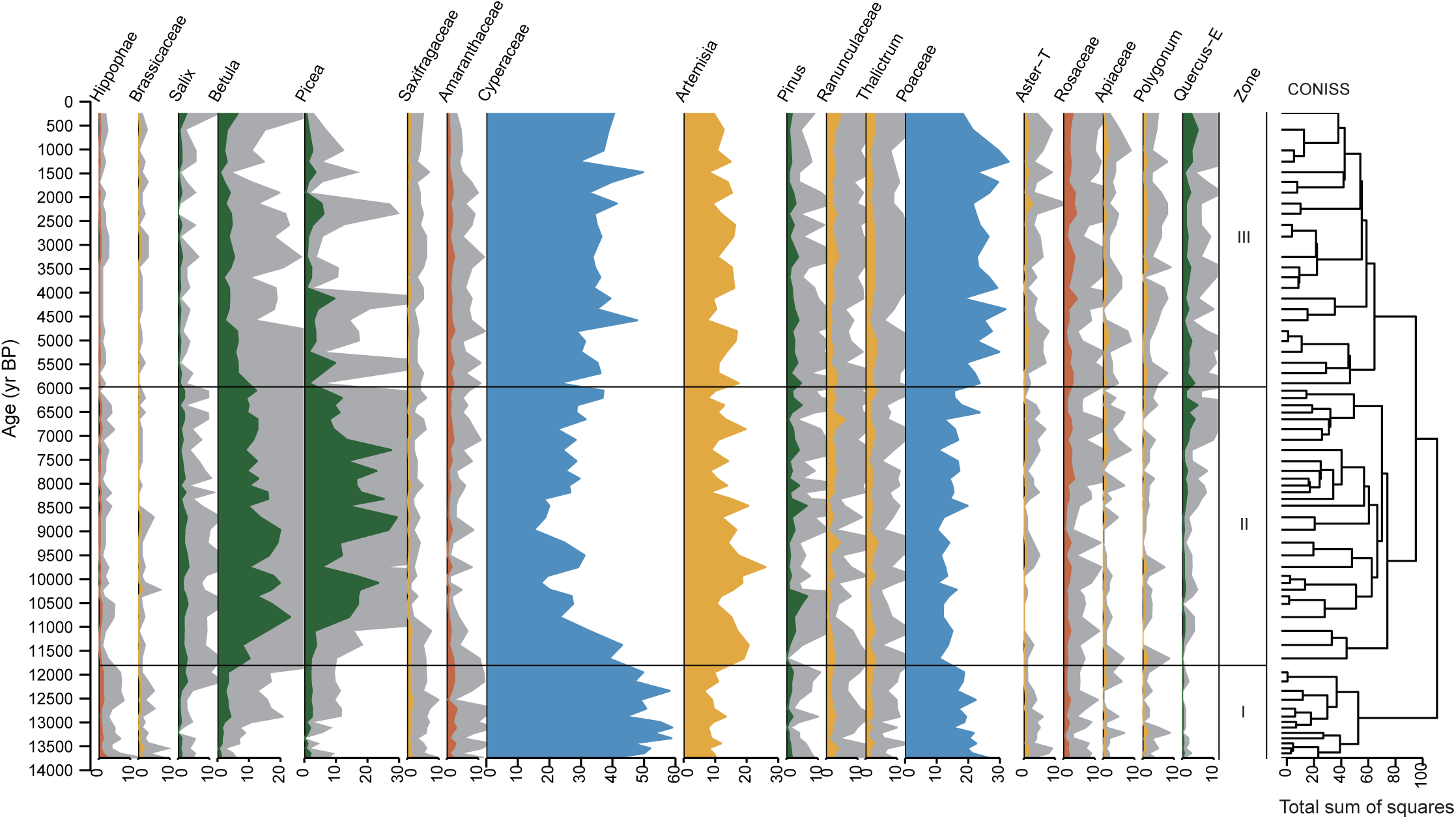
Relative abundance of terrestrial pollen taxa over the past 14 ka from the Zhagaer Co sediment core. Green denotes trees, brown shrubs, yellow forbs, and blue graminoids. CONISS-defined zones are indicated on the right. Grey shading is a five-fold exaggeration for better visualization.

### 4.4 RDA of sedaDNA community composition along the Picea gradient

RDA indicated a significant association between plant community composition and *Picea* abundance (permutation test, P < 0.05; Figure 5). The constrained axis (RDA1) explained 5.4% of the total variance, and the overall model was significant based on 999 permutations. A parallel analysis using pollen-derived *Picea* as the constraining variable yielded highly comparable results in terms of variance explained and model significance (Supplementary Note 2).

**Figure 5.**
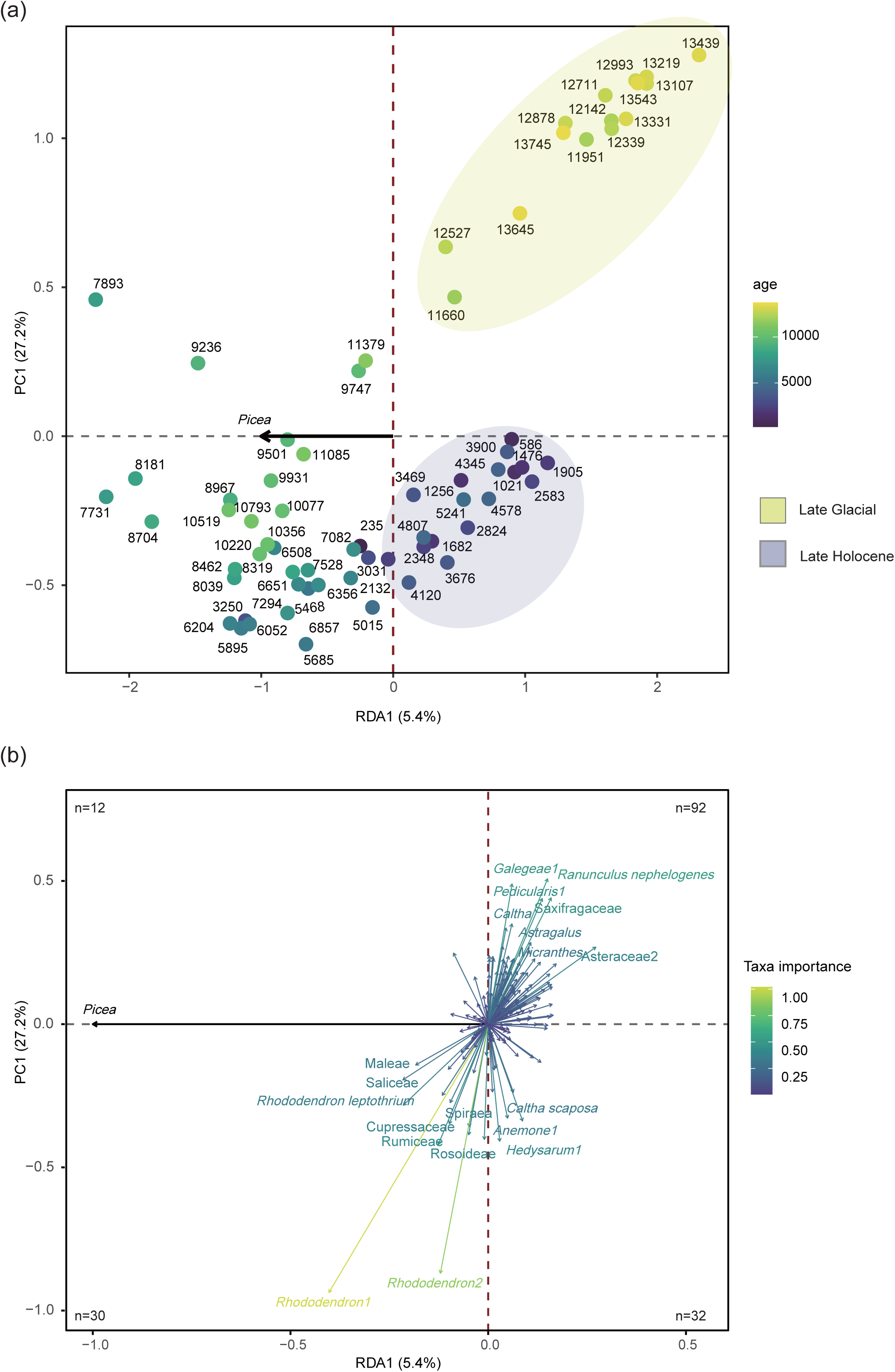
Redundancy analysis (RDA) of terrestrial plant community composition constrained by *Picea* abundance in the Zhagaer Co sedaDNA record. (a) Distribution of sample scores along the Picea-constrained forest gradient. (b) Taxon scores showing associations with forest-dominated versus open-alpine conditions. The number of taxa in each quadrant is indicated.

Samples distributed along the RDA1 axis showed clear temporal differentiation (Figure 5a). Early to mid-Holocene samples were associated with negative RDA1 values, whereas late Glacial and late Holocene samples were generally associated with positive RDA1 values. Late Glacial samples were primarily located in the first quadrant, while late Holocene samples occurred more frequently in the fourth quadrant, although they were not restricted to this quadrant. Taxa positively associated with *Picea* were primarily woody plants, including *Betula*, *Alnus*, *Rhododendron*, and Saliceae, whereas negatively associated taxa were mainly alpine and open-habitat groups such as *Ranunculus nephelogenes*, *Galegee*, *Pedicularis*, *Hedysarum*, *Anemone*, and *Caltha scaposa* (Figure 5b). Samples and taxa positioned along the negative direction of RDA1 represent the “winner” end of the gradient, whereas those on the positive side constitute the “loser” end used in the subsequent contrasts.

Based on the “winner–loser” classification, family-level comparisons revealed consistent differences in functional composition between the two groups, with winners tending to be dominated by woody taxa (e.g., Ericaceae, Salicaceae, and Betulaceae), whereas losers were more often represented by herbaceous taxa, particularly alpine and open-habitat families such as Papaveraceae, Saxifragaceae, and Poaceae (Figure 6a). Within the subset of loser samples, clear compositional differences were observed between the late Glacial and late Holocene intervals (Figure 6b). Although both groups occupied the same side of the RDA1 gradient, their family-level composition differed. Late Glacial samples showed higher representation of Gentianaceae, Phrymaceae, Poaceae, Rosaceae, and Cyperaceae, whereas late Holocene samples were characterized by relatively greater contributions of Primulaceae, Fabaceae, Orobanchaceae, Saxifragaceae, Ranunculaceae, and Papaveraceae. These patterns indicate compositional differentiation within open-alpine assemblages across cold phases.

**Figure 6.**
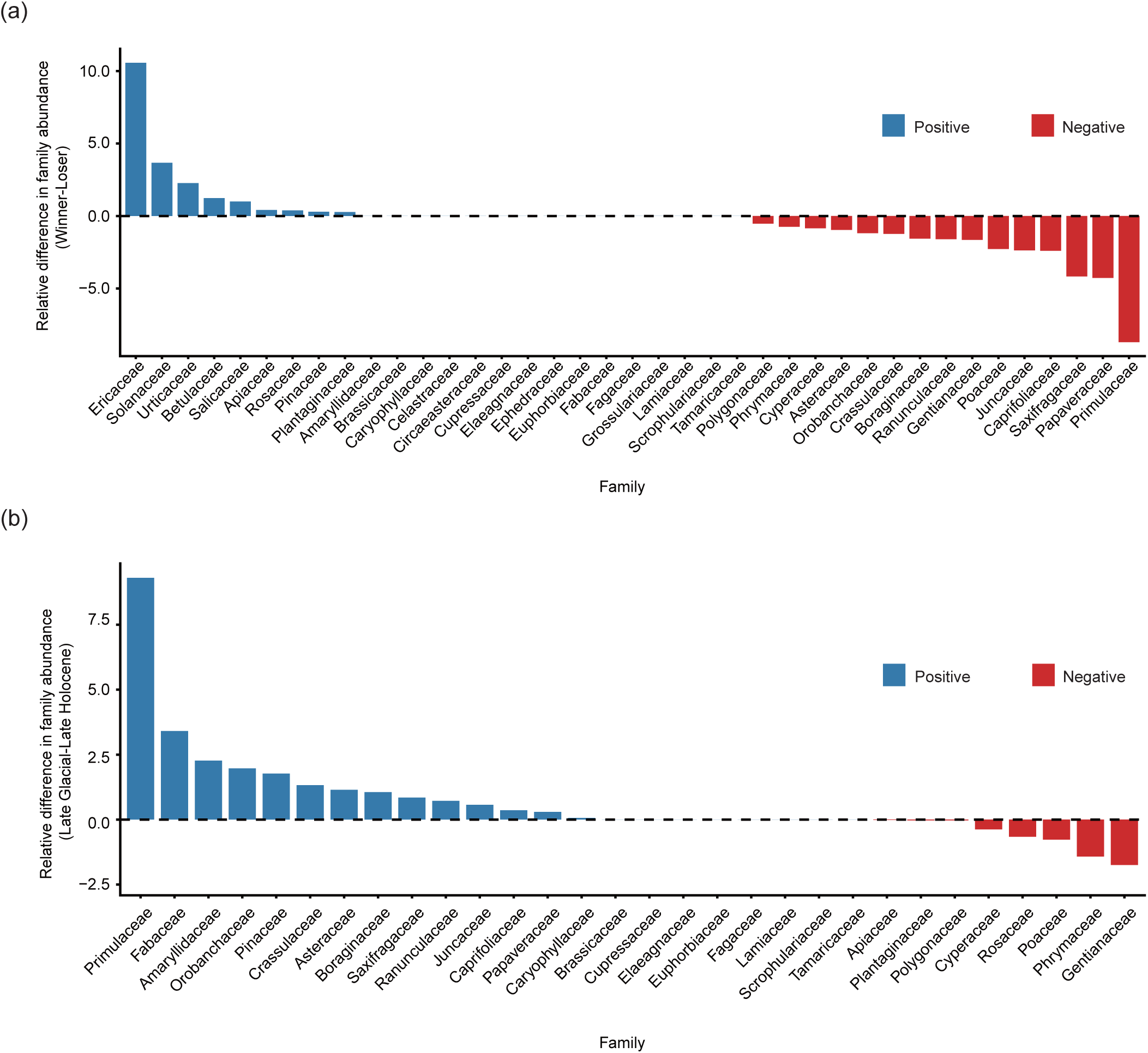
Family-level differences in relative abundance associated with RDA1-defined winner-loser groups and their comparison between Late Glacial and Late Holocene phases. (a) Family-level differences in relative abundance between RDA1-defined winner and loser groups. (b) Family-level differences in relative abundance between the late Glacial and late Holocene within the RDA1-defined loser group.

## 5. Discussion

### 5.1 Complementary ecological signals in sedaDNA and pollen

Both sedaDNA and pollen delineate three broadly consistent vegetation phases in our core, with comparable temporal boundaries separating a late Glacial alpine-dominated phase, an early to mid-Holocene interval marked by expanded forest cover, and a late Holocene phase characterized by reduced tree cover and partial alpine reopening (Figure 3; Figure 4). This consistency in temporal patterns indicates that both proxies capture the major transitions in vegetation composition at Zhagaer Co over the past 14 ka. Despite this overall agreement, the two proxies differ in taxonomic richness, taxonomic resolution, and spatial representativeness. These differences are particularly evident in taxonomic richness and taxonomic resolution. As shown in Table 2, the sedaDNA record recovered substantially more taxa and achieved considerably finer taxonomic assignments, with nearly half of all identifications resolved to species level, whereas pollen identifications were largely restricted to genus and family levels. This difference highlights the contrasting strengths of the two proxies and demonstrates the greater resolving power of sedaDNA for reconstructing vegetation composition (Jørgensen et al. 2012; Liu et al., 2020). In addition, the two proxies differ in their spatial representativeness. In our record, pollen consistently shows higher proportions of arboreal taxa during the early to mid-Holocene, like *Picea*, *Betula*, and *Pinus*, than sedaDNA, whereas sedaDNA records a relatively stronger contribution of herbaceous and alpine taxa (Figure 3; Figure 4). These contrasts likely reflect differences in source area and taphonomic processes, with sedaDNA predominantly capturing vegetation from the lakeshore and local catchment (Alsos et al., 2018; Li et al., 2021), whereas pollen integrates signals over a broader source area through wind dispersal, often involving long-distance transport, including extra-regional forest stands (Li et al., 2020; Zhang et al., 2017). These differences influence the type and scale of vegetation information captured by each proxy. As a result, pollen and sedaDNA do not represent independent or conflicting signals, but rather provide complementary perspectives across spatial and taxonomic scales.

**Table 2.** Summary of the taxonomic resolution of the sedaDNA and pollen approaches.

| Methods | Number of samples | Number of taxa | Taxonomic resolution (%) |  |  |  |  |  |  |
| --- | --- | --- | --- | --- | --- | --- | --- | --- | --- |
|  |  |  | Family | Subfamily | Tribe | Subtribe | Genus | Species | Clade |
| sedaDNA | 68 | 513 | 4.09 | 2.34 | 6.82 | 3.7 | 34.1 | 48.3 | 0.585 |
| Pollen | 67 | 41 | 41.5 | 0 | 0 | 0 | 58.5 | 0 | 0 |

The observed differences could be explained by methodological and taphonomic constraints inherent to the two proxies. Methodologically, each proxy is subject to distinct limitations. Pollen identification is constrained by morphological similarity among grains, which groups multiple taxa into broad pollen types, whereas sedaDNA resolution depends on barcode markers and the completeness of reference databases, which may limit taxonomic assignment for certain lineages (Taberlet et al., 2007). Beyond methodological constraints, taphonomic processes may also contribute to differences between the two proxies. Pollen grains are efficiently dispersed by wind, allowing signals from multiple spatial sources to be integrated, and high pollen production in many taxa further enhances their representation in pollen assemblages (Li et al., 2022). In contrast, sedaDNA is delivered to lake sediments through water-mediated pathways and is therefore strongly dependent on hydrological connectivity, resulting in a more localised signal. In addition, sedaDNA is more susceptible to degradation, which further influences its representation in the sediment record (Parducci et al., 2017; Giguet-Covex et al., 2019; Jia et al., 2021).

### 5.2 Comparative vegetation dynamics across lake records in the Nianbaoyuze Mountains

Our sedaDNA and pollen analyses (Figure 3; Figure 4) from Zhagaer Co in the southeastern Nianbaoyuze Mountains show broadly consistent patterns of vegetation change through time. Vegetation around this region was dominated by open alpine meadow communities during the late Glacial, followed by an increasing forest cover during the early to mid-Holocene and vegetation shifted again towards open communities dominated by shrub and alpine meadow during the late Holocene. Similar regional vegetation dynamics have been reported across the Nianbaoyuze Mountains and the eastern Tibetan Plateau, despite variability in taxonomic composition among sites.

During the late Glacial (13.7 – ∼11.6 cal ka BP), both sedaDNA and pollen records from Zhagaer Co indicate that vegetation was dominated by alpine meadow communities (Figure 3; Figure 4), reflecting open and largely treeless conditions. This is further supported by the presence of characteristic high-elevation taxa in the sedaDNA record, which today occur primarily above the modern treeline (Wu et al., 1994–2013), consistent with predominantly above-treeline vegetation under cold climatic conditions. At the regional scale, pollen records from the Nianbaoyuze Mountains and the surrounding eastern Tibetan Plateau, primarily from high-elevation sites, similarly document the dominance of alpine meadow taxa and very low arboreal pollen percentages during this period (Schlütz & Lehmkuhl, 2009; Kramer et al., 2010b; Herzschuh et al., 2014; Li et al., 2023). This consistent pattern across multiple sites indicates that vegetation was relatively homogeneous and predominantly above the treeline across the region during the late Glacial. Together, these records suggest that late Glacial vegetation across the region was primarily structured by strong climatic constraints, especially low temperatures and a shortened growing season, which maintained open alpine communities and limited tree establishment. A slight increase in *Picea* and *Betula* around ∼13 ka (Figure 4) may indicate the initial response of woody taxa to improving climatic conditions, although overall arboreal representation remains low.

During the early to mid-Holocene (∼11.6–5.5 cal ka BP), vegetation at Zhagaer Co underwent a major ecological transition from open alpine meadow communities to increasing woody cover, as consistently indicated by both sedaDNA and pollen records (Figure 3; Figure 4). Graminoids and alpine forbs declined, while woody taxa, particularly *Picea*, *Betula*, *Rhododendron*, and Salicaceae, increased markedly, indicating a shift towards more forested conditions and an upslope advance of the treeline during this period (Xu et al., 2025). At the regional scale, similar increases in woody taxa are observed across the Nianbaoyuze Mountains and the eastern Tibetan Plateau (Kramer et al., 2010a; Herzschuh et al., 2014; Li et al., 2023), suggesting a broadly coherent pattern of forest expansion. However, differences in the relative representation of woody taxa are apparent among sites. At Zhagaer Co, *Picea* appears to be one of the dominant woody taxa, as indicated by both sedaDNA and pollen records, although *Betula* is also relatively abundant in the pollen assemblages. A similar co-occurrence of *Picea* and *Betula* is observed in nearby records such as Emu Co (Li et al., 2023). In contrast, pollen records from other lakes in the region, including Ximen Co and Naleng, are characterised by a stronger dominance of *Betula* (Kramer et al., 2010a; Herzschuh et al., 2014). These differences in the relative representation of woody taxa indicate that vegetation patterns were not spatially uniform, even at a sub-regional scale. Such spatial variability raises the question of what processes controlled the local dominance of different tree taxa. The relatively high abundance of *Picea* at Zhagaer Co may reflect local environmental conditions that favour its persistence, as *Picea* is still present in the region today. In addition, relict populations of *Picea* have been reported near Mageng Co, located only ∼20 km from Emu Co in the Nianbaoyuze Mountains (Schlütz and Lehmkuhl, 2009; Li et al., 2023), indicating the presence of nearby seed sources within the region. Such local availability of *Picea* may have facilitated its establishment and expansion in certain sites. Together, these observations suggest that, although forest expansion occurred widely across the region, the dominance of tree taxa was influenced by local seed sources and dispersal limitations, which may also help explain the stronger dominance of *Betula* at sites such as Ximen Co.

During the late Holocene (since ∼5.5 cal ka BP), vegetation around Zhagaer Co shifted towards more open alpine conditions, characterised by the expansion of alpine meadow and shrubland communities, suggesting a transition towards cooler and drier conditions. This shift is supported by both sedaDNA and pollen records (Figure 3; Figure 4), which show increasing abundances of graminoids and forbs together with shrubs such as *Rhododendron* and Rosaceae, accompanied by a decline in tree cover. Similar changes have been reported across multiple sites in the eastern Tibetan Plateau (Herzschuh et al., 2014; Li et al., 2023), indicating a broadly coherent regional pattern of forest retreat during the late Holocene. Notably, several records also show a slight increase in *Quercus* pollen during this period, suggesting a consistent regional signal. This increase may reflect enhanced input from broadleaved vegetation at lower elevations through long-distance dispersal. However, although *Quercus* has been associated with relatively dry conditions in previous studies, its ecological significance in high-elevation records remains uncertain. Taken together, these observations suggest that late Holocene vegetation changes were primarily driven by regional climatic forcing, while additional factors such as local environmental conditions, human activity, and atmospheric CO₂ likely modulated vegetation dynamics.

### 5.3 Forest expansion as a potential ecological filter shaping “winner-loser” dynamics since the late Glacial

Forest expansion appears to structure plant assemblages along the *Picea*-defined gradient. In the RDA ordination, taxa associated with increasing woody cover are clearly separated from those linked to open and alpine habitats (Figure 5b), indicating a directional shift in community composition rather than a neutral reshuffling. These patterns are consistently observed across taxonomic levels and suggest the emergence of contrasting responses among taxa. Such responses can be interpreted as “winner–loser” dynamics, with some lineages increasing in association with woody expansion while others decline under the same conditions.

Woody shrubs and forest-margin taxa tend to emerge as ecological “winners” during phases of forest expansion at Zhagaer Co. Genera typical of treeline and forest-edge environments, including *Rhododendron*, *Betula*, *Alnus*, and members of Salicaceae, cluster toward the forest-associated (high-*Picea*) end of the gradient and generally exhibit consistently positive winner-loser scores (Figure 5b; Table S1). This pattern is also evident at the family level, where Ericaceae, Salicaceae, Betulaceae, and Rosaceae show similarly positive values (Figure 6a). Together, these taxon- and family-level results indicate that forest expansion tends to favour woody lineages associated with treeline and forest-margin environments. The coherence between sample-level separation and taxon-level responses suggests that forest expansion may have influenced community composition in a directional manner. Rather than affecting taxa randomly, increasing forest cover appears to favour woody lineages, while disadvantaging light-demanding alpine herbs and graminoids. This pattern is consistent with ecological filtering associated with increasing woody cover, whereby more shaded and mesic conditions at forest margins may promote woody establishment in treeline systems (Greiser et al., 2024; Chen et al., 2024). Stage-dependent shrub–tree interactions documented in modern treeline systems—initial facilitation of tree seedlings by shrubs followed by increasing competition for light as woody cover develops—may provide a plausible explanation for this pattern (Callaway, 1998; Germino et al., 2002). Notably, *Rhododendron* shows a delayed increase relative to other woody taxa and becomes abundant only when *Picea* expands upslope (Figure 3), which may reflect greater edaphic requirements and the development of mesic, organic-rich soils beneath established conifer stands (Bharali et al., 2014).

In contrast, forest expansion is associated with reduced representation of alpine and open-habitat taxa along the same forest–alpine gradient. In our RDA results, alpine forbs and graminoids such as *Pedicularis*, *Caltha*, *Ranunculus* and *Astragalus* cluster toward the low-*Picea* end of the ordination (Figure 5b; Table S1), and sample-group contrasts identify Saxifragaceae, Papaveraceae, Caryophyllaceae, Asteraceae, Poaceae and Cyperaceae as consistently lower-scoring families (Figure 6a). These results indicate that alpine and open-habitat lineages are consistently disadvantaged across taxonomic levels. Their decline under increasing woody cover likely reflects reduced light availability and the progressive loss of open microsites as forest structure develops, conditions that disadvantage cold-adapted, light-demanding taxa during shrub and tree encroachment at forest margins (Liu et al., 2023). Many of these lineages share functional traits such as low stature and specialisation to exposed alpine habitats, making them poorly suited to shaded or structurally complex environments. Several of these groups also exhibit long-term alpine niche conservatism, having diversified and persisted in high-elevation or open biomes throughout the late Cenozoic (Sun et al., 2017). The consistency of loser identities across regions (Liu et al., 2021; Shen et al., 2025) further supports this interpretation. Comparisons with modern forest–alpine systems, where increasing woody cover promotes shade-tolerant shrubs but reduces light-demanding herbs and graminoids, reinforce this interpretation (Majasalmi et al., 2020; Liu et al., 2023). Together, these results indicate that forest expansion consistently disadvantages light-demanding, open-habitat lineages, reflecting a directional filtering process that selectively excludes alpine taxa.

### 5.4 Asymmetric cold-open assemblages during late Glacial and late Holocene

The late Glacial and late Holocene alpine phases represent non-equivalent ecological states, despite both being characterised by relatively cold climatic conditions. During the late Glacial, cold conditions supported an open alpine landscape dominated by cold-adapted alpine taxa, such as Saxifragaceae, Orobanchaceae, and Papaveraceae (e.g. *Saxifraga*, *Pedicularis*, *Meconopsis*) (Figure 5b; Figure 6b; Table S1). In the late Holocene, although cooling also led to the reappearance of some alpine taxa (e.g. Saxifragaceae), the overall community composition was markedly different, with increased contributions from graminoids and meadow taxa, including Cyperaceae, Poaceae, and Rosaceae (*Carex*, *Pooideae*, *Sibbaldia*) (Figure 5b; Figure 6b; Table S1). This indicates that comparably cold climatic conditions did not produce equivalent ecological outcomes, suggesting that climate alone could not fully account for the observed differences in community composition. These differences likely reflect contrasting ecological trajectories between the two periods. The late Holocene alpine communities developed following extensive forest expansion during the early to mid-Holocene and subsequent forest retreat, which likely left lasting imprints on community composition, resulting in assemblages that may reflect both past forest presence and current climatic conditions (e.g. Herzschuh, 2020). In addition, atmospheric CO2 concentrations during the late Glacial (∼190 ppm) were substantially lower than during the late Holocene (∼280 ppm), which may have further constrained plant productivity and tree establishment, particularly at high elevations where CO2 partial pressure is already reduced (Herzschuh et al., 2011).

Human pastoral activity may also have acted as an additional driver shaping late Holocene alpine assemblages. The increased abundance of grazing-tolerant graminoids (Poaceae, particularly Pooideae, *Poa* and *Koeleria*), together with wet-meadow and disturbance-tolerant sedges (Cyperaceae, e.g. *Carex* subgen. *Carex*, *C.* subgen. *Euthyceras*, *Carex przewalskii*), as well as meadow taxa such as *Sibbaldia* and *Potentilla*, is consistent with vegetation types commonly associated with grazed alpine meadows (Figure 6b; Table S1; Schlütz & Lehmkuhl, 2009; Miehe et al., 2019). Independent palaeoecological and archaeological evidence suggests that pastoralism was established on the eastern Tibetan Plateau during the late Holocene (e.g. Schlütz & Lehmkuhl, 2009; Chen et al., 2015; Liu et al., 2024); However, these taxa are not uniquely indicative of human activity and their relative importance could not be clearly distinguished from climatic and ecological processes. Together, these results suggest that differences between late Glacial and late Holocene alpine communities were shaped by a combination of climatic forcing and historical contingency, with climate determining overall alpine conditions and forest legacy influencing the specific composition of plant assemblages, and with potential contributions from human activity.

## 6. Conclusion

Our integrated sedaDNA and pollen records from Zhagaer Co indicate that Holocene vegetation dynamics on the eastern Tibetan Plateau were closely associated with forest expansion and retreat along the forest–alpine ecotone. Forest expansion during the early to mid-Holocene appears to have favoured woody forest-margin taxa while reducing alpine forbs and graminoids, giving rise to consistent “winner–loser” patterns across taxonomic levels. These results suggest that shifts in forest cover were associated with directional changes in alpine plant assemblages, consistent with patterns expected under ecological filtering rather than neutral turnover.

Comparison of the late Glacial and late Holocene cold-open phases further shows that alpine communities that re-emerged after Holocene forest retreat did not fully reassemble into a Late Glacial-like state. Instead, the late Holocene assemblage contains a greater contribution of meadow and shrub taxa, suggesting that post-forest alpine communities followed a path-dependent trajectory influenced by the ecological legacy of prior forest expansion.

Together, these findings highlight the importance of treeline history in shaping alpine vegetation states over millennial timescales. While climatic forcing likely initiated forest expansion and retreat, late Holocene pastoral activity may have contributed to the persistence of open alpine meadow–shrub systems. Our results suggest that alpine vegetation patterns on the eastern Tibetan Plateau reflect the combined influence of climate-driven treeline dynamics, ecological filtering, and human land use, emphasising the role of ecological legacies and anthropogenic influences in long-term vegetation change.

## Author Contributions

All authors have made substantial contributions to the manuscript. U.H. and Y.Z. designed the study. U.H., X.C. and Y.Z. led the interpretation and writing. Y.Z. wrote the first draft of the manuscript. X.C. contributed to the core collection and Y.Z. performed DNA lab work. S.C. performed the pollen lab work. Y.Z., K.R.S., and F. T. implemented the data analysis and statistical analyses. All authors discussed the results and provided intellectual input to the manuscript.

## Declaration of competing interest

The authors declare that they have no known competing financial interests or personal relationships that could have appeared to influence the work reported in this paper.

## Data availability

The raw NGS sequencing data are deposited in the European Nucleotide Archive (ENA) at EMBL-EBI under the accession PRJEB98242 (Zhagaer Co).

## Acknowledgements

We thank Kai Yi for help with sub-sampling, Janine Klimke and Jasmin Tauchelt for their technical support in the paleogenetic laboratories. This study was supported by the National Natural Science Foundation of China (Grant No. 42471179), the Sino-German Mobility Programme (grant no. M-0359) and the China Scholarship Council (CSC, Grant No. 202204910035 to Y.Z.).

## Competing Interests statement

The authors declare no competing interests.

## Supplementary Materials

**Table S1**. Taxa assigned to the four quadrants of the RDA ordination, ranked by their contributions (species scores) along the RDA axes.

**Supplementary Note 1: Quality control of Zhagaer Co sedaDNA results**

**Supplementary Note 2: Evaluation of laboratory controls**

## Notes

### Competing Interest Statement

The authors have declared no competing interest.

