## Supplementary Information for "Forest expansion and open vegetation responses during the past 14 ka at Zhagaer Co on the eastern Tibetan Plateau"

By Zhang et al.

This file includes:

Supplementary table:

Table S1

Supplementary Notes:

Supplementary Note 1: Quality control of Zhagaer Co sedaDNA results

Supplementary Note 2: Evaluation of laboratory controls

Table S1. Taxa assigned to the four quadrants of the RDA ordination, ranked by their contributions (species scores) along the RDA axes.

| **Quadrant Q1 (92)** | | | **Quadrant Q2 (12)** | **Quadrant Q3 (30)** | **Quadrant Q4 (32)** |
| --- | --- | --- | --- | --- | --- |
| *Ranunculus nephelogenes* | *Ranunculus bungei* | *Eritrichium* | *Alnus cremastogyne* | *Rhododendron1* | *Hedysarum1* |
| Galegeae1 | *Draba* | *Anaphalis* | *Parnassia1* | *Rhododendron2* | *Anemone1* |
| Saxifragaceae | *Micranthes divaricata* | *Pedicularis18* | *Chrysosplenium* | Rumiceae | *Caltha scaposa* |
| *Pedicularis1* | Apioideae | Boraginaceae1 | *Nicotiana tabacum* | Rosoideae | Swertiinae |
| Asteraceae2 | *Lonicera* | *Bistorta macrophylla* | *Betula* | Cupressaceae | *Thalictrum* |
| *Caltha* | Asteraceae | *Gentiana* | Euclidieae | Spiraea | *Sibbaldia* |
| *Astragalus* | *Rheum* | *Carex przewalskii* | *Hippophae tibetana* | *Rhododendron* *leptothrium* | *Carex subgen Euthyceras1* |
| *Micranthes* | *Pedicularis20* | *Pedicularis22* | *Veronica subgen Stenocarpon* | Saliceae | *Paraquilegia* |
| *Rhodiola2* | *Saxifraga* | Androsace | Potentilleae2 | Maleae | Fagaceae |
| *Caltha gracilis* | *Eremogone* | *Potentilla* | *Ephedra* | Apioideae5 | Cichorieae |
| *Ranunculus1* | *Delphinium spirocentrum* | *Pedicularis12* | *Circaeaster agrestis* | *Saussurea* | *Corydalis* |
| *Meconopsis3* | Poeae | Galegeae | *Trigonotis* | *Ribes1* | Anthemideae |
| *Pedicularis anas* | *Ranunculus nephelogenes* var *longicaulis* | *Rhodiola* |  | *Callitriche palustris* | *Koenigia* |
| *Saxifraga sinomontana* | *Melanoseris violifolia* | *Pedicularis24* |  | *Cardamine macrophylla* | *Delphinium* |
| *Pedicularis11* | *Koenigia1* | *Chrysosplenium axillare* |  | *Saussurea mucronulata* | *Silene* |
| *Ranunculus rufosepalus* | Polygonoideae | Asteroideae3 |  | Spiraeeae | *Carex subgen Carex* |
| *Primula1* | *Pedicularis15* | *Carex alatauensis* |  | Apioideae7 | *Euphorbia sect Helioscopia* |
| Asteroideae1 | *Ligularia tsangchanensis* | *Aconitum tanguticum* |  | *Elsholtzia densa* | *Lonicera1* |
| Tussilagininae | *Indocypraea montana* | *Youngia* |  | *Urtica3* | *Melica scabrosa* |
| *Saussurea dzeurensis* | *Corydalis crispa* | *Pinus* |  | IRL clade | *Sibbaldia cuneata* |
| *Rumex* | *Allium* | Triticodae |  | Apioideae2 | *Rubus sachalinensis* |
| *Pedicularis17* | Juncaceae | Myricaria1 |  | *Urtica2* | *Corydalis conspersa* |
| *Ranunculus2* | Tussilagininae1 | Asteroideae2 |  | Artemisiinae1 | *Koeleria* |

Table S1 (continued)

| **Quadrant Q1 (92)** | | | **Quadrant Q2 (12)** | **Quadrant Q3 (30)** | **Quadrant Q4 (32)** |
| --- | --- | --- | --- | --- | --- |
| Ranunculoideae | *Meconopsis* | *Carex myosuroides* |  | *Carex subgen Carex1* | *Juniperus microsperma* |
| *Lagotis2* | *Syncalathium chrysocephalum* | *Allium1* |  | *Saxifraga3* | Pooideae |
| *Meconopsis integrifolia* | Asteroideae | Colurieae |  | *Limosella aquatica* | *Vicia sativa* |
| Gnaphalieae | Astereae | *Hippophae1* |  | Potentilleae | BOP clade |
| Apioideae6 | *Saxifraga1* | Scrophulariaceae |  | *Knorringia sibirica* | *Eutrema* |
| *Pedicularis2* | Pooideae1 | Delphinieae1 |  | Carduinae | *Ajuga* |
| *Faberia pinnatifida* | *Rhodiola chrysanthemifolia* | *Salvia incertae sedis1* |  | *Thermopsis alpina* | *Bistorta paleacea* |
| Asteraceae1 | Poeae2 |  |  |  | *Lancea tibetica1* |
|  |  |  |  |  | Poeae3 |

**Supplementary Note 1: Quality control of Zhagaer Co sedaDNA results**

In total, 550 ASVs from Zhagaer Co sedaDNA sequencing matched sequences in the "TP_db" database at 100% similarity. Non-metric multidimensional scaling (NMDS) analysis of the three independent PCR replicates from each sediment sample revealed high compositional consistency among replicates, except for two top-layer sample. This likely reflects increased micro-scale heterogeneity and patchy deposition in surface sediments, where multiple DNA sources and recent inputs may lead to less homogeneous community signals (Figure S1).

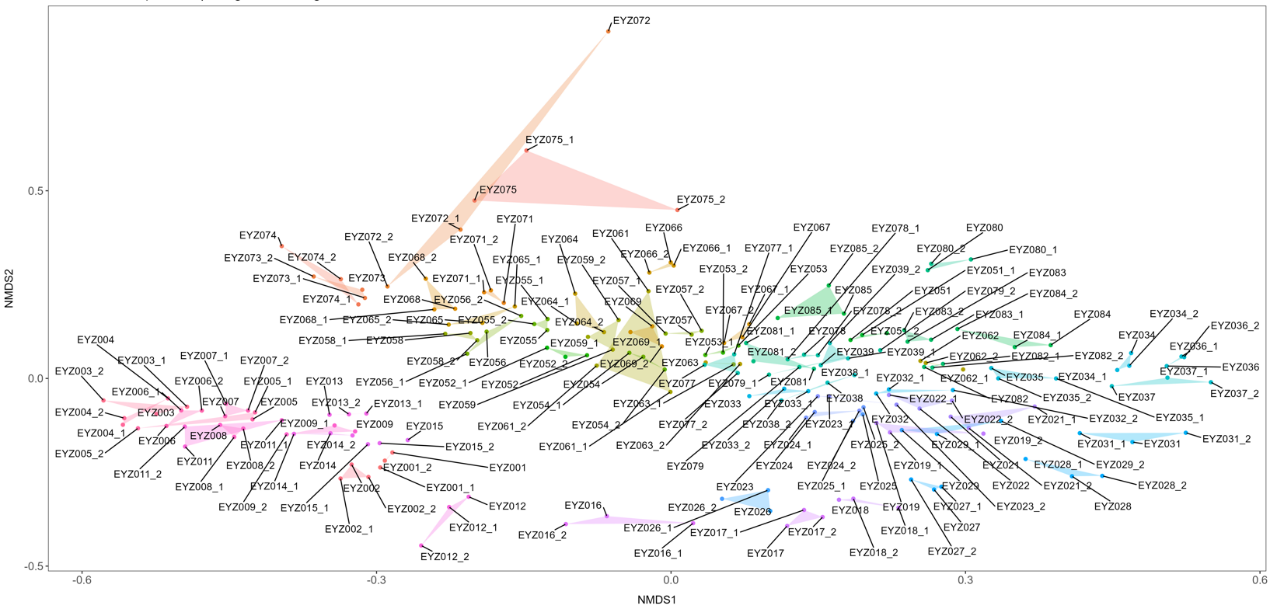

Figure S1. Non-metric multidimensional scaling (NMDS) results. Labels in the legend indicate the composite laboratory number, depth (cm), and the number of PCR replicates of the sample. For example, ‘EYZ001_1’ means this sample is the second PCR replicated at a depth of 673 cm.

**Supplementary Note 2: Evaluation of laboratory controls**

***Sequencing summary and control assessment***

The dataset AMPG-77 contains in total 46642730 read counts and 550 sequence types derived from totally 270 sample replicates (3 PCR replicates for each sediment sample) and 66 negative control replicates (47 extraction controls (3 PCR replicates for the extraction control of each extraction batch, only extraction chemicals) and 19 PCR negative controls (only PCR chemicals)). The total read count in the sample PCR replicates is 32128984 (69 % from total) and the total read count in all negative control replicates is 14513584 (31 % from total), whereof only 162 (0.0003% from total) are detected in PCR negative controls with PCR chemicals only. 33861 read counts (0.007 % from total) that were attributed to four sequence types only occurring in the controls, but not in the samples, were discarded from the dataset.

The negative controls of the extractions show basically two main sequence types (*Picea* and *Pinus*) summarizing to 5489845 read counts (38% from total read count in all negative control replicates) in 37 extraction controls. Most likely these sequence types are a result of chemical contamination of the DNA extraction Kit. However, the contamination in the controls has only a minor effect on the sample compositional signal, because there is (1) a high replicability of PCR replicates in the samples (see Figure S3), (2) a very similar temporal in *Picea* confirmed by an independent pollen record of Zhagaer Co (R=0.577, p<0.001; Figure S1) and (3) a high degree of concordance among RDA ordinations constrained by *Picea* abundance from independent sedaDNA and pollen datasets, as indicated by Procrustes analysis (Figure S2, Figure S3, table S2). Based on our evaluation, we kept all sequence types detected in the sample dataset.

***Independent DNA extraction and metabarcoding dataset***

The independent metabarcoding analysis followed basically the same procedure, but sediments were extracted with the PowerSoil Kit and a starting volume of about 300mg per sample. Extractions were done in replicates and extracted DNA was pooled and concentrated via GeneJet PCR purification. PCR and pooling was performed as described in the Material & Methods section. The raw data of the independent metabarcoding analysis is publicly available under the ENA BioProject accession number: PRJEB98242.

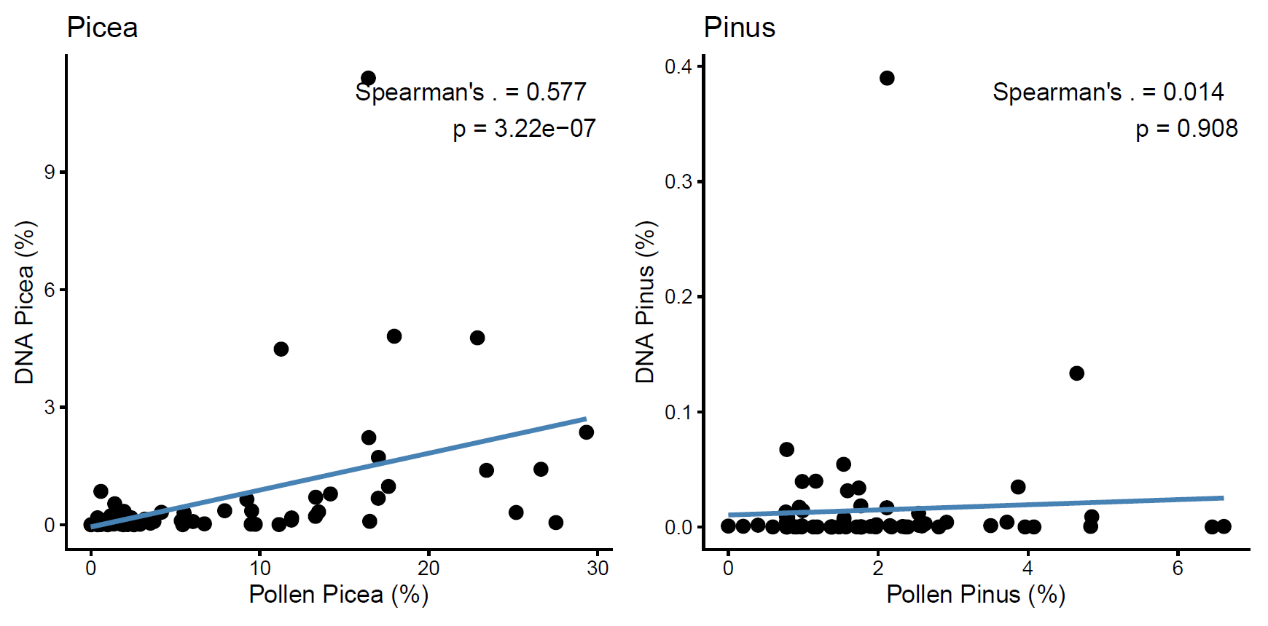

Figure S1. Relationship between pollen and sedaDNA *Picea* (%)

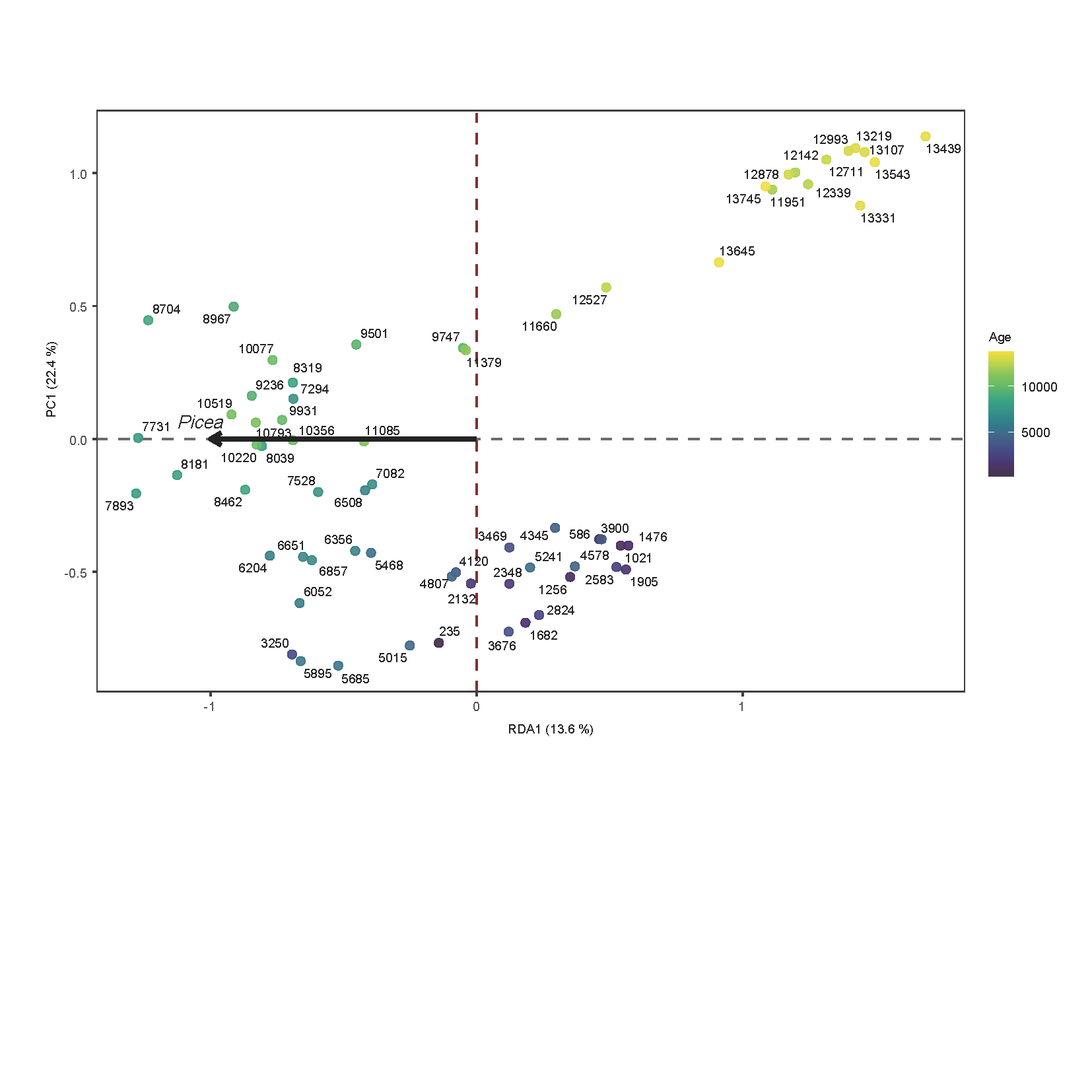
Figure S2 Redundancy analysis (RDA) of sedaDNA-inferred plant community composition constrained by *Picea* pollen abundance, used as a proxy for forest cover. Each point represents a sample and is coloured by age.

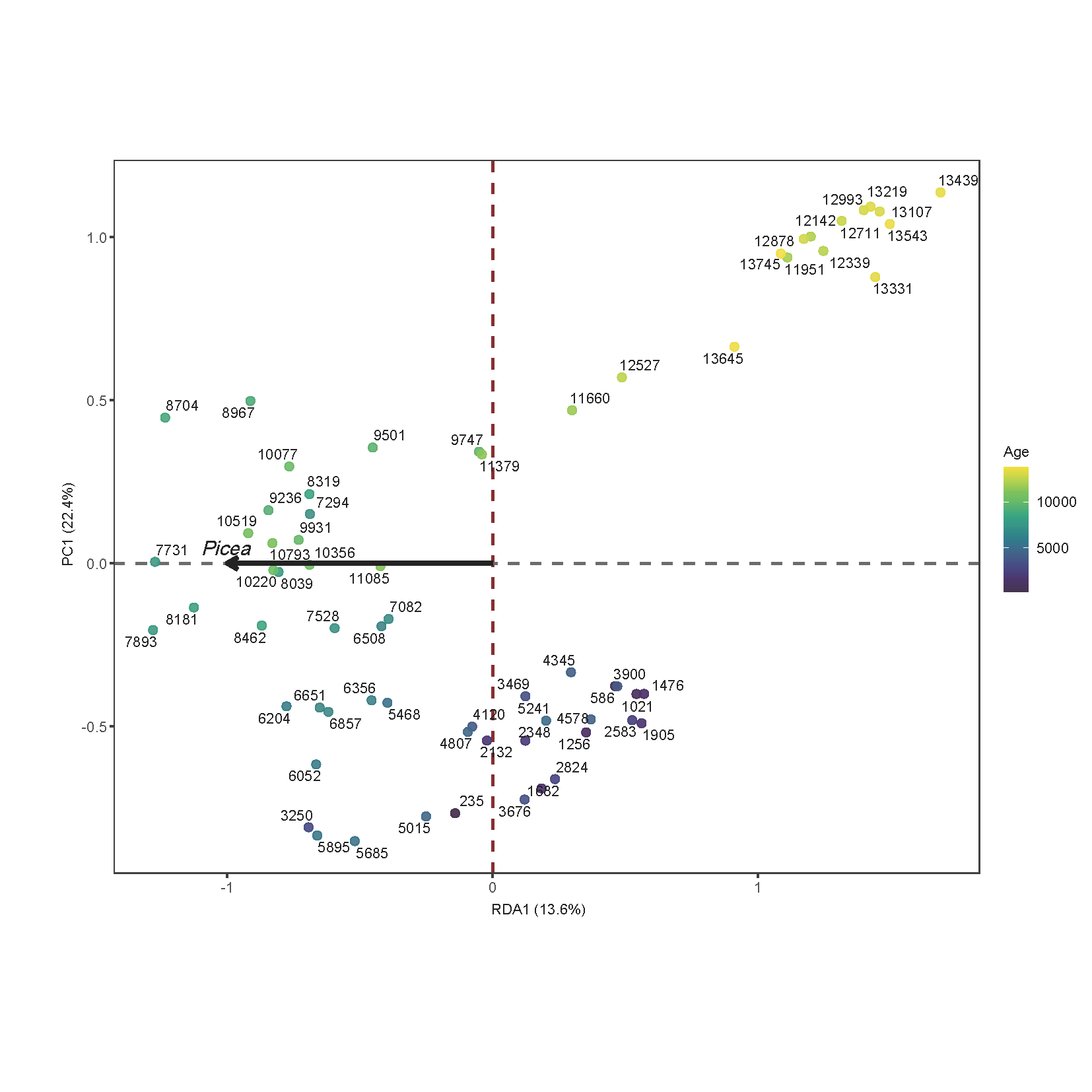

Figure S3 Redundancy analysis (RDA) of sedaDNA-inferred plant community composition constrained by *Picea* New sedaDNA abundance, used as a proxy for forest cover. Each point represents a sample and is coloured by age.

***Consistency of ordinations across datasets***

To evaluate whether the *Picea* signal in the sedaDNA dataset remains a reliable indicator of forest dynamics despite potential contamination, we performed a cross-proxy validation based on constrained ordination. Redundancy analysis (RDA) was applied to the sedaDNA community matrix using *Picea* abundance as the constraining variable. To assess the robustness of the resulting ordination, the same community matrix was constrained separately using *Picea* abundance derived from three sources: the original sedaDNA dataset, an independent sedaDNA dataset generated from a pollen dataset (Figure S2) and a separate DNA extraction (Figure S3). This design allowed us to evaluate whether different representations of the forest indicator produced comparable ecological gradients.

The resulting RDA site scores (first two axes) were compared using Procrustes analysis, and statistical significance was assessed using the protest permutation test (999 permutations) implemented in the vegan package in R. Only samples with matched ages across datasets were retained for comparison.

The Procrustes analyses revealed a high degree of concordance among the three RDA ordinations (Table S2). Pairwise comparisons yielded consistently high correlations (r > 0.89, p = 0.001), indicating that the primary community gradient associated with *Picea* is broadly consistent across proxies. Concordance was highest between the two sedaDNA-based ordinations, while slightly lower similarity was observed between pollen- and DNA-constrained ordinations.

**Table S2. Procrustes correlations among RDA ordinations constrained by different *Picea* proxies. Values represent Procrustes correlation coefficients, with associated p-values in parentheses.**

|  | **Original sedaDNA** | **Pollen** | **Independent sedaDNA dataset** |
| --- | --- | --- | --- |
| **original sedaDNA** | — | 0.942(p = 0.001) | 0.942(p = 0.001) |
| **Pollen** | 0.942(p = 0.001) | — | 0.893(p = 0.001) |
| **independent sedaDNA dataset** | 0.942(p = 0.001) | 0.893(p = 0.001) | — |
